# Genetic, metabolic, and molecular insights into the diverse outcomes of diet-induced obesity in mice

**DOI:** 10.1101/2021.09.02.458729

**Authors:** Alexis Maximilien Bachmann, Jean-David Morel, Gaby El Alam, Sandra Rodríguez-López, Tanes Imamura de lima, Ludger J.E. Goeminne, Giorgia Benegiamo, Maroun Bou Sleiman, Johan Auwerx

## Abstract

Overweight and obesity are increasingly common public health issues worldwide, leading to a wide range of diseases from metabolic syndrome to steatohepatitis and cardiovascular diseases. While the increase in the prevalence of obesity is partly attributable to changes in lifestyle (i.e. increased sedentarity and changes in eating behaviour), the metabolic and clinical impacts of these obesogenic conditions varies between sexes and genetic backgrounds. The conception of personalised treatments of obesity and its complications require a thorough understanding of the diversity of responses to conditions such as high-fat diet intake. By analysing nine genetically diverse mouse strains, we show that much like humans, mice respond to high-fat diet in a genetic- and sex-dependent manner. Physiological and molecular responses to high-fat diet are associated with expression of genes involved in immunity and mitochondrial function. Finally, we find that mitochondrial function may explain part of the diversity of physiological responses. By exploring the complex interactions between genetics and metabolic phenotypes via gene expression and molecular traits, we shed light on the importance of genetic background and sex in determining metabolic outcomes. In addition to providing the community with an extensive resource for optimizing future experiments, our work serves as an exemplary design for more generalizable translational studies.

## Introduction

Today, 15% of the worldwide population is obese, and the prevalence of the disease is increasing steadily, driven by changes in lifestyle (diet, physical activity, and stress). Obesity is often observed in conjunction with metabolic syndrome, type-2 diabetes mellitus^1^ and non-alcoholic fatty liver disease (NAFLD)^2^. Frequent symptoms include insulin resistance, hypertension and dyslipidaemia, which constitute risk factors for cardiovascular diseases, cancer, and diabetes, making these three related diseases the leading cause of mortality and premature disability^3^. Although obesity, diabetes and fatty liver often overlap, patients may present any combination of them. It is commonly understood that the wide variety of responses to obesity are determined by environmental and genetic factors. A study that gave identical diets to a large human cohort found highly variable glycemic responses to identical meals^4^. While genome-wide association studies in humans have met some success in finding variants causing extreme cases of obesity, the genetic determinants of more common forms of obesity remain elusive^5^. This is partly because of the difficulty in estimating environmental factors, and because external measurements of body weight or BMI are not sufficient to distinguish between different types of obesity^5^.

Another approach for understanding the diverse aetiology of obesity is the use of animals, especially mouse models, in which one can easily control genetic and environmental factors^6^. The simplest of these models are animals fed with a fat-supplemented diet (high-fat diet, HFD). A caveat of these studies is often the focus on a single mouse strain, such as the C57BL/6J reference strain, sometimes genetically modified, failing to model the diversity of outcomes present in the human population. Health benefits of precision dietetics in mice are highly strain-dependent^7^, similar to what is observed for rodent studies on drug therapies^8^ or toxicology^9,10^. Findings originating from a single strain and sex not only translate poorly to humans but also to other mouse strains^11^. A recent study highlighted how a large genetically diverse population of mice could model the diversity and interindividual variability in non-alcoholic fatty liver disease (NAFLD), upon feeding with a high-sucrose and -fat diet and find susceptible and resistant mouse strains^12^. More generally, we hypothesize that findings that hold true in different mouse strains and subspecies are much more likely to also be conserved in humans^13^.

To address this, we systematically characterized the diversity and extent of the physiological, molecular, and functional changes induced by HFD across diverse *Mus musculus* inbred strains, subspecies, and sexes. While we observe some expected responses to the diet, most metabolic and molecular parameters are affected by HFD in a manner that is dependent on sex and strain. Our observations indicate that the genetic repertoire of inbred laboratory mouse strains encodes a wide range of metabolic responses to diet-induced obesity (DIO), modelling different levels of metabolic disease severity or associated comorbidities. All metabolic traits, mitochondrial activity and transcriptome data collected in this study can be explored with an online, interactive interface at https://lisp-lms.shinyapps.io/Project_CC_Founder_App/. This resource enables researchers to quickly examine the diverse manifestations of DIO in mice and pick an appropriate mouse model for individual obesity-associated diseases.

## Results

### Phenotypic responses to HFD are dependent on mouse genetic backgrounds and sexes

10 female and 10 male animals from 9 inbred mouse strains were fed with either a chow diet (CD) or high-fat diet (HFD) from the age of 8 to 21 weeks and various metabolic and fitness phenotypes were measured (Figure 1A). Eight of these strains are known as the founders of the Collaborative Cross (CC)^14^ and Diversity Outbred (DO) populations^15^. The DBA/2J was added as a ninth strain to leverage our laboratories’ expertise with the BXD recombinant inbred lines^16,17^ which are generated from crosses of the C57BL/6J and DBA/2J strains. Three distinct *Mus musculus* subspecies are represented in this panel: *Mus musculus musculus* (PWK/PhJ), *Mus musculus castaneous* (CAST/EiJ) and *Mus musculus domesticus* (all other strains including the wild-derived WSB/EiJ)^18^.

**Figure 1:**
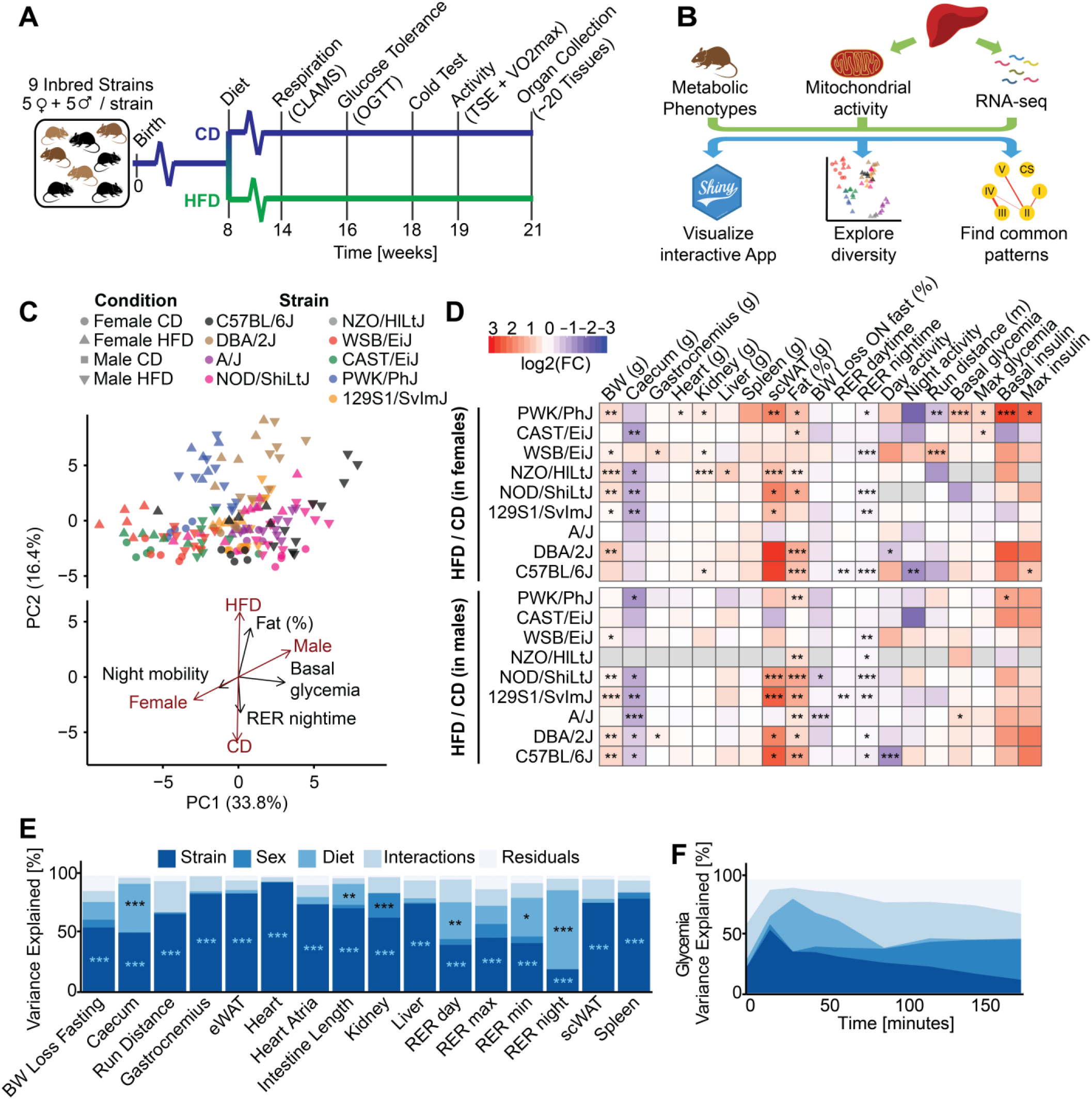
Variation in metabolic traits is driven by distinct contributions of genetic background, sex, and diet. A) Scheme of the phenotyping pipeline of the study. 9 mouse inbred strains were tested (C57BL/6J, DBA/2J, A/J, 129S1/SvImJ, NOD/ShiLtJ, NZO/HILtJ, WSB/EiJ, CAST/EiJ and PWK/PhJ). 5 females or 5 males per diet (CD or HFD) were measured per strain. Five mice per condition were fed an HFD from 8 weeks of age to 21 weeks with CD matched animals. Respiration, tolerance to glucose and cold, exercise capacity and spontaneous activity were tested. Mice were sacrificed at 21 weeks of age after an overnight fast and organ collection was performed. B) Schematic of the liver molecular phenotyping and data analysis. Livers were used for RNA-Seq and measurements of mitochondrial activity. C) Principal component analysis plot of the metabolic and clinical traits measured in the study. Individual data are shown (top). Loadings are shown for selected phenotypes as well as gender and diet (bottom). Principal component 1 is associated with a combination of strain and sex, while PC2 is associated with the diet. Black arrows: phenotypic traits. Brown arrows: covariates. D) Heatmap of the effects of diet on a selection of measured metabolic traits, in males or females. *P<0.05, **P<0.01, ***P<0.001, t-test (HFD vs CD), corrected for multiple testing with the Benjamin-Hochberg false discovery rate (BH-FDR) procedure. E) Proportion of variance of metabolic traits explained by a linear combination of strain, sex and diet effects, their interactions and the residuals of the linear model (Phenotype ∼ Strain * Sex * Diet), expressed as the mean type II sums of squares divided by the total mean type II sum of squares. Mean sum of squares is equal to the sum of squares divided by the corresponding degrees of freedom. *P<0.05, **P<0.01, ***P<0.001, ANOVA test corrected for multiple testing. F) Variance of glycemia explained by strain, sex and diet effects, their interactions during oral glucose gavage test (OGTT) over time.

We measured metabolic traits and liver-centered molecular traits to assess the conserved or sex- and strain-specific effects of HFD as well as correlations between biological and phenotypic layers (Figure 1B). As a multivariate approach, we used principal component analysis (PCA) to determine each trait varied in the different conditions (Figure 1C). Principal component 1 (PC1, 33.8 % variance explained) is associated with a combination of sex and strain. Whereas principal component 2 (PC2, 16% variance explained) segregated mice based on diet. Some phenotypes, such as the fat percentage and respiratory exchange ratio (RER), closely correlated with diet, whereas others, like basal glycemia and activity were more associated with strain and sex. The covariance components of diet and sex partially overlap because body weight and organ weights are higher in both males and HFD-fed animals. By using a multivariate approach, we determined that sex and strain dominate variation in phenotypic traits, followed by HFD.

Next, we examined the effects of HFD on phenotypes in males and females of each strain (Figure 1D). The response to diet was highly variable between strains and sexes. The A/J and CAST strains gained limited weight upon feeding a HFD (Figure 1D, Figure S1A). Subcutaneous white adipose tissue (scWAT) weight increased in most strains, except the WSB/EiJ and CAST/EiJ. WSB/EiJ and CAST/EiJ are wild-derived strains known for their resistance to obesity and high percentage of lean mass^19^. Female PWK/PhJ mice had the most comprehensive response to HFD (e.g. glucose and insulin levels, RER during the night) (Figure 1D). Altogether, our data highlight that the responses to HFD are sex- and strain-specific without a conserved signature across metabolic parameters.

Of note, weight gain and decrease of RER in response to HFD did not always accompany glucose intolerance (Figure S1A, Figure S1B & Figure S1C, online interface). For example, the C57BL/6J, DBA/2J and PWK/PhJ similarly gained weight upon HFD but disruption of glycemia was much greater in the C57BL/6J. This is particularly interesting because some strains present an uncoupling of often-associated metabolic disorders including obesity, diabetes and metabolic substrate utilisation. In addition, low body temperature during cold tolerance tests was associated with glucose intolerance in most strains (Figure S1D), as is also the case in humans^20^. The lowest temperatures were observed in C57BL/6J, NOD/ShiLtJ and NZO/HILtJ, which also reached the highest glycaemia during the oral glucose tolerance test (OGTT). The A/J and NZO/HILtJ mice had much lower maximum running distance during the VO_2_max experiment (Figure S1E & Figure S1G), perhaps because their RER peaked immediately, showing quick saturation with CO_2_, while other strain’s RER increased gradually (Figure S1E). Body weight loss upon overnight fasting, an indicator of metabolic flexibility^21^, was consistently lower in HFD-fed mice across strains and sexes (Figure S1F).

To quantify the relative contributions of the different biological factors to metabolic outcomes, we computed the phenotypic variance explained by each factor of the study using ANOVA and type II sum of squares (Figure 1E). The effect of the diet on the RER was larger during the night than during the day (Figure 1E), reflecting the main feeding and activity period for mice. While dynamic parameters (e.g. RER), were diet-dependent, morphological measurements, such as organ weights, were mainly strain-dependent, with the exception of caecum weight, which is known to be highly affected by the diet^22^. Unexpectedly, strain effects explained most of the variation of adipose tissue mass (eWAT and scWAT) highlighting the influence of genetics on fat storage. The effects on glycemia during OGTT were particularly striking (Figure 1F). While basal glycemia (glucose 0 min) was influenced to a large extent by strain, sex and diet interactions; the maximal glycemia (reached at 15 minutes) was predominantly a function of strain, reflecting genetic differences in glucose absorption. Finally, the values at 30, 45 and 60 minutes were much more influenced by diet, as the speed of glucose return to basal levels (clearance) is greatly influenced by diabetic-like phenotypes that arise in some strains under HFD. Altogether, these data highlight the complex interactions between strain, sex and diet for various metabolic traits and show that the spectrum of variation of metabolic traits in mice partially reflects the spectrum of healthy and pathologic conditions found in humans with an obesogenic diet.

### The liver transcriptome shows a context-dependent diet signature

To link variation of metabolic traits with molecular data, we collected and analysed liver gene expression by RNA-Seq in 3 mice per condition, sex, and strain. Like phenotype-based multivariate analyses, liver gene expression clustered samples mainly according to strain and sex (Figure 2A).

**Figure 2:**
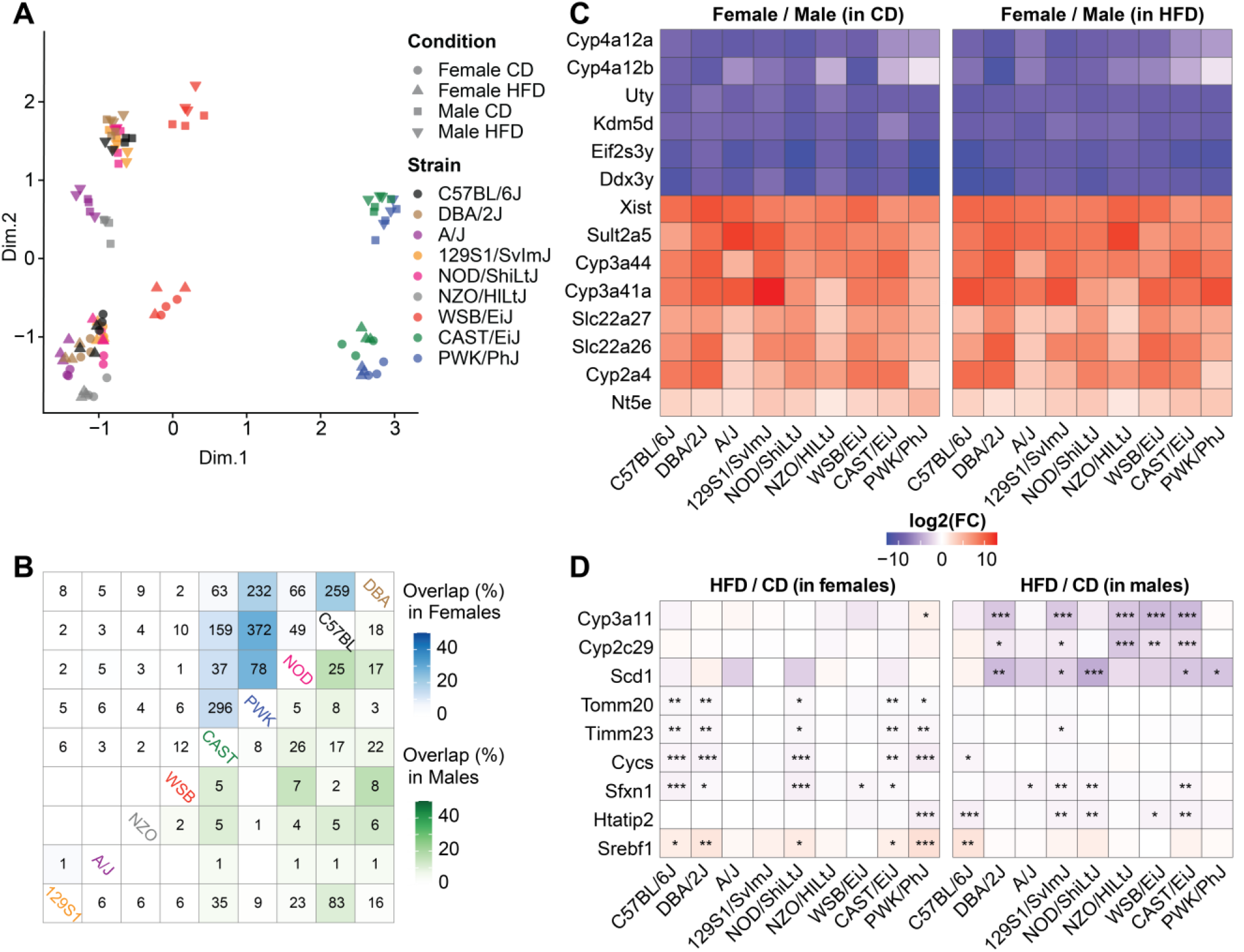
The transcriptome signature of HFD differ between strains and sexes. A) Multidimensional scaling (MDS) plot of liver RNA-Seq separates animals by strain and sex. Euclidean distances between each animal were calculated pairwise based on the leading log_2_-fold changes for the 500 most differentially expressed genes. Each dot or triangle represents a single female or male mouse respectively. Each color is assigned to a specific strain. B) Number of overlapping differentially expressed genes (Diet effect; HFD/CD) across strains of the same sex (female: top left, male: bottom right). Color code representing the percentage of overlapping genes is based on the Jaccard index. C) Heatmaps of log_2_ fold change of the 14 most conserved sexually dimorphic genes across strains and diets. All the fold changes are significant with BH-FDR adjusted p-value<0.05. D) Heatmaps of log_2_ fold change of the 9 genes affected by diet in the highest number of strains at the same time. The same color scale was used for panel C & D. *P<0.05, **P<0.01, ***P<0.001, adjusted p-value corrected for multiple testing with the BH-FDR procedure.

The effects of the diet could be seen on a strain by strain-basis but were far weaker than sex effects (Figure S2A). This contrasted with the PCA of the phenotypes, where the effect of diet was much more pronounced, and strains and sexes overlapped (Figure 1C). While the effect of sex was conserved in both diets (Figure S2B, top), the diet effects were different between sexes (Figure S2B, bottom). To quantify differences/similarities between the transcriptomic profiles of the mice, we performed Pearson’s correlations for each pairwise combination between samples and these comparisons were categorised according to the strain, sex and diet similarities (Figure S2C). Pairwise transcriptome correlations consistently followed genetic relatedness, with correlations decreasing from same strain > same subspecies > different subspecies. Having the same diet only increased correlations when the strain and sex were also the same (Figure S2C, comparison on the right), showing that transcriptomic diet effects are only reproducible within a given strain and sex. The number of overlapping differentially expressed genes (DEGs) of HFD vs. CD-fed animals across genetic backgrounds contrasted across sexes (Figure 2B). In females, C57BL/6J, DBA/2J, CAST/EiJ and PWK/PhJ mice showed the highest number of overlapping DEGs, whereas in males this was apparent mostly in 129S1/SvImJ and C57BL/6J. For example, PWK and C57BL/6J had 372 DEG in common in females and only 8 in males, whereas C57BL/6J and 129S1/SvImJ had only 2 DEG in common in females and 83 in males. Therefore, differences and similarities between strains and between diets are not symmetrical between females and males. This means that the use of male or female mice needs to be considered with caution while choosing a particular strain to model a metabolic process. Conversely, sex differences are much better conserved across strains and diets (Figure S2D). Since sex differences were exceptionally well conserved across mouse strains, we asked if they would also reflect the sexual dimorphism in the human liver transcriptome. We compared the expression fold change between sexes of mouse genes to that of their human orthologs in the Genotype-Tissue Expression’s (GTEx) database of human liver transcriptomes (Figure S3). The overall correlation was significantly positive, although small, indicating an overall trend of conserved sex effects. The genes with highest fold changes were sexually dimorphic in both mice and humans, but while some of them (e.g. *UTY* and *CIB3*) varied in the same direction, others (*SULT1A1, EIF2S3L*) were sexually dimorphic in both species, yet inverted.

### HFD affects immunity and protein translation in a sex- and strain-dependent manner

While individual diet-dependent genes differ across strains and sexes, we investigated whether those different genes might still participate in some of the same biological pathways. Rather than analyse every combination of strain, sex or diet separately, we used the overall variance explained by each parameter and performed gene set-enrichment analysis (GSEA) to understand which pathways are driven by each covariate (Schematic of the method, Figure 3A). As expected, HFD had consistent effects on lipid and carbohydrate metabolism (Figure 3B). Interestingly, sex, diet and their interactions all affected immune processes. At the cellular component level (GOCC), mitochondrial content and secretory vesicles were strongly affected by sex and diet, while ribosomes (ribonucleoprotein complex) were affected by sex:diet interactions. At the transcription factor level, the target genes of Lymphoid Enhancer Binding Factor 1 (*Lef1*) were particularly affected by sex, whereas those of Estrogen Related Receptor 1 (*Err1*) are affected by diet^23^.

**Figure 3:**
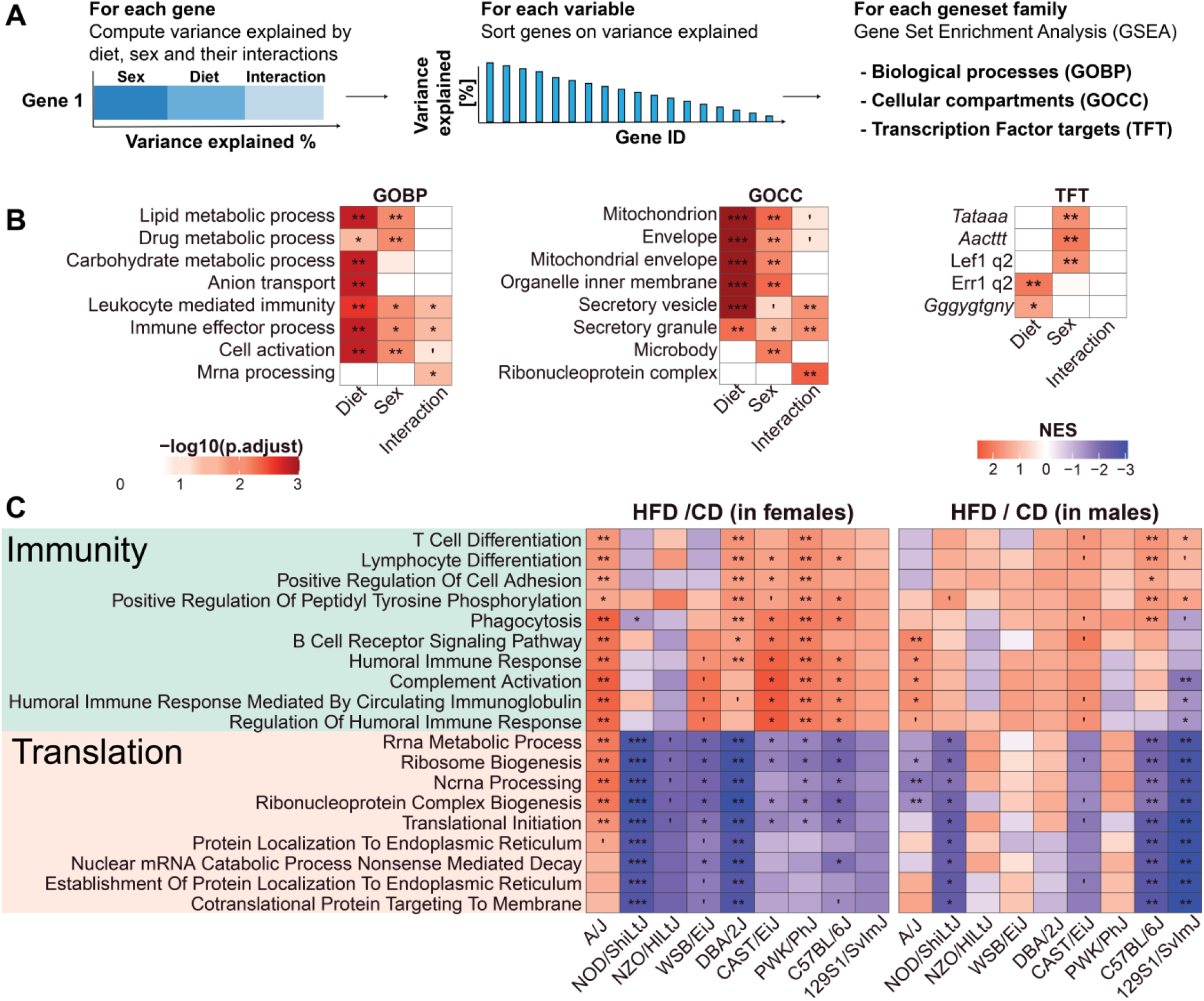
HFD affects immunity, lipid metabolism and liver translation in a sex and strain-dependant manner. A) Scheme of geneset enrichment analysis based on variance explained by diet, sex and their interaction. (1) For each transcript, we computed the variance explained based on a linear model of strain, sex, diet and their interactions (Transcript ∼ Strain + Diet + Sex + Diet:Sex). Then the percentage of variance explained of each transcript was assigned to specific parameters (Diet, Sex, and their interactions) (2) for ranking of the genes and then (3) to perform geneset enrichment analysis for each parameter. B) Results of geneset enrichment analysis based on variance explained by diet, sex and their interaction. From left to right each enrichment was performed on gene ontology of biological processes (GOBP), cellular component (GOCC) and transcription factors. Percentage of variance explained was measured using *variantPartition* package. C) Enrichment analysis of the diet effect on liver gene expression separated by sex and strain. Gene set enrichment analysis was performed on the sign(log_2_(FC)) * –log_10_(adjusted p-value) per strain per sex. The selected gene sets add the highest score for the global response to the diet in a specific sex.

To highlight the differences between strains, we performed another GSEA of the effect of diet on each strain/sex combination and displayed the most conserved ones (Figure 3C). Enriched gene sets clustered into two main categories: immunity and translation/mRNA processing. Overall, adaptive immune responses were enriched upon HFD to a higher extent in females compared to males (Figure 3C). This reflects sex-specific manifestations of inflammation and immune response in the context of fatty liver^24^. Genesets associated with translation, endoplasmic reticulum (ER) and mRNA processing were reduced upon HFD in most strains, perhaps reflecting an activation of the integrated stress response^25^. In humans, the integrated stress response is activated in obese and/or patients with fatty liver^26^. Although these pathways were the most conserved in the population, there were still important differences between strains. Indeed, A/J females showed a significant positive enrichment of translation associated gene sets, opposite to the other strains (Figure 3C).

### Liver mitochondrial activity *in vivo* correlates with metabolic traits

Due to the documented importance of mitochondria as sensor and effector of metabolism, we measured the activities of citrate synthase (CS), the main pace-making enzyme of the tricarboxylic acid cycle (TCA) and the different complexes of the Electron Transport Chain (ETC; Complex I-V) in liver tissue. CS activity was used here as a readout of entry of fuel into the TCA cycle and as a proxy of mitochondrial content^27^. Citrate synthase itself was strongly affected by strain, with the lowest activity present in A/J mice (Figure 4A), which has a known high-impact genetic variant in the coding sequence of the gene encoding citrate synthase itself^28–31^. We also observed a consistent reduction of Complex I and Complex III activity upon HFD in males only (Figure 4B). In addition, we computed the variation explained by strain, sex, diet and their interactions on CS activity and on each complex activity (Figure 4C). It is striking to observe the extent to which mitochondrial complexes of the ETC are differently affected by biological variables. Indeed, while Complex I activity is highly strain-dependent, variation of Complex II activity is predominantly explained by sex differences. Most of the variation of Complex III activity can be attributed to the variation of Citrate Synthase activity (Figure 4C).

**Figure 4:**
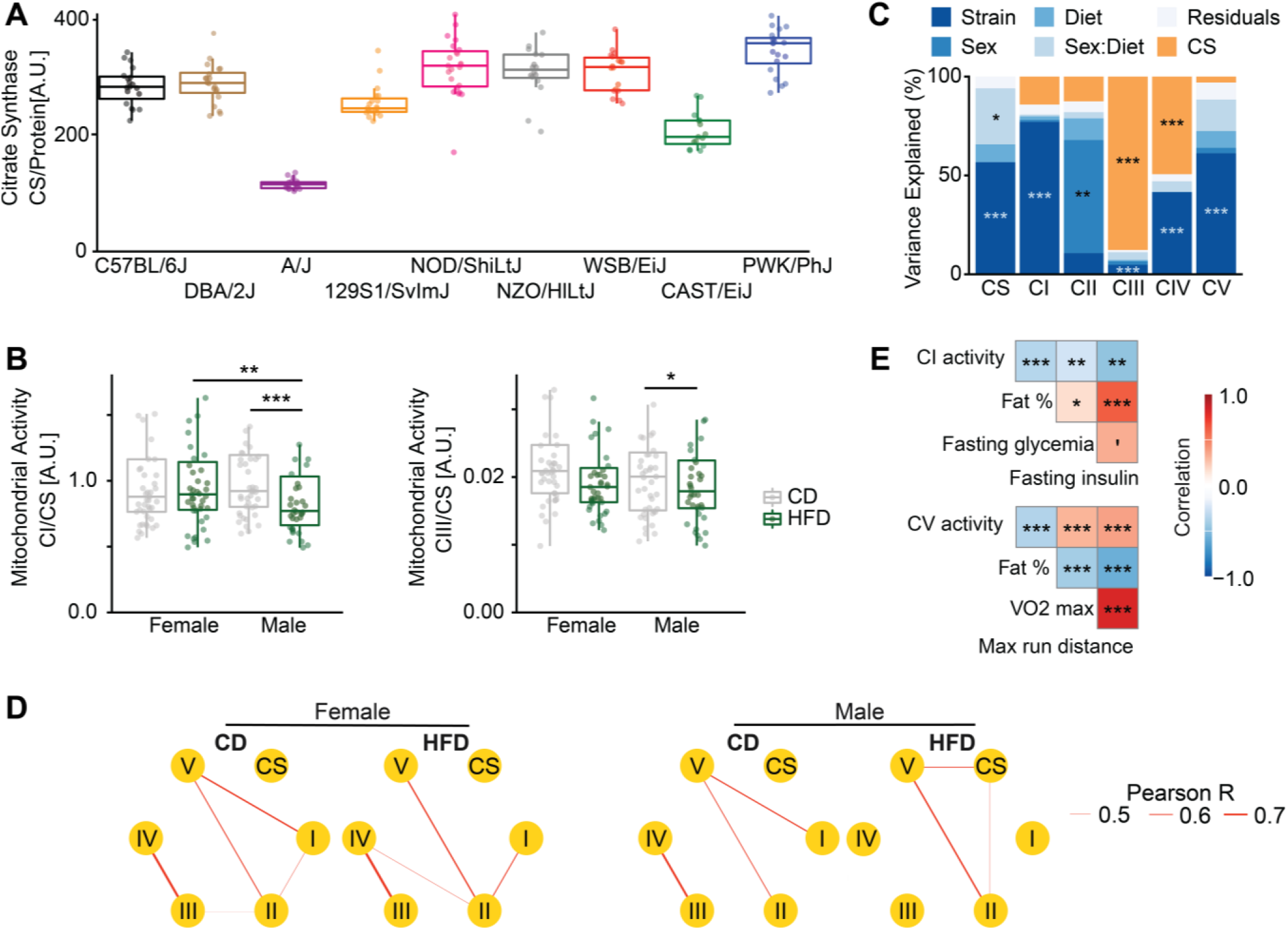
Liver mitochondrial activity correlates with metabolic phenotypes, and correlations between complexes is disrupted by HFD in males. A) Citrate synthase activity (A.U.) is principally affected by strain. B) Complex II and Complex III activity (normalised by citrate synthase activity) is affected by diet in sex-specific manner. C) Proportion of variance of liver mitochondrial activity explained by strain, sex, diet, sex-by-diet interactions (sex:diet) and citrate synthase (CS) activity. Each bar corresponds to the activity of the different complexes of the electron transport chain as well as the citrate synthase activity or a model which takes all the complexes together into account (ALL). *P<0.05, **P<0.01, ***P<0.001, ANOVA test. D) Correlation network of liver mitochondrial complex activities in each condition. The width of the edges corresponds to the magnitude of positive correlation between complexes. Only significant edges are shown (adjusted-p <0.05). E) Spearman’s correlations between the activity of liver mitochondrial complexes and essential metabolic traits. *P<0.05, **P<0.01, ***P<0.001, correlation test, corrected for multiple testing with the BH-FDR procedure.

We then computed the correlation between ETC complexes for each experimental condition (Figure 4D). Complex II and V activity are consistently correlated across conditions, whereas correlation between Complex I and Complex V activity was lost upon HFD. In addition, correlations of complex I and complex II activity were sex-specific, present in females and not in males. Finally, complex III and complex IV activity associations are not present in males fed HFD, which highlights the potential difference of the response to HFD in males and females (Figure 4D). Complexes I and V activity both negatively correlated with fat percentage, suggesting that strong mitochondrial activity is protective against weight gain or that HFD affects negatively mitochondrial activity. Complex I activity further correlated negatively with fasting glycemia and insulin, whereas complex V activity positively correlated with VO2 max and running distance, indicators of long-term stamina (Figure 4E). Thus, the activity of mitochondrial complexes I and V, the main entry and exit points of mitochondrial electron transport, may reflect the type of response to obesogenic diet. In the future, the correlation map (Figure 4D) of the activity of mitochondrial complexes, could be used as a type of metabolic “fingerprint”, predicting metabolic responses, although this warrants further investigation.

## Discussion

Here, we presented a detailed phenotypic and liver transcriptomic analysis, as well as mitochondrial activity of 9 mouse inbred strains of both sexes and with two different diets (CD and HFD). Our focus was to understand which traits and phenotypic layers show the most consistent or the most strain-dependant responses to sex and diet. Our data suggest that traits that are better conserved across mouse strains have a greater likelihood of being translatable to humans. When looking at morphological phenotypes, such as tissue weights, we found that the genetic effect of the strain predominated over any other (Figure 1E). The body weight and fat percentage, which we would expect to be primarily driven by environmental factors, were mostly a function of the strain, highlighting the importance of genetics in all morphological traits, including fat tissues. Conversely, functional phenotypes representing metabolism or physical performance, such as RER, maximum run distance, or the glycemic response were much more affected by the diet and interactions between the diet, strain and sex. In the OGTT test, it was striking that while maximum glycemia was mostly a function of the strain, likely reflecting the speed of glucose absorption, the glucose clearance was more influenced by diet and diet/sex/strain interactions (Figure 1F). The speed of return to normal glycemia is a feature of metabolic syndrome and diabetes, and it seems that its alteration by diet is partially conserved across strains.

In the liver transcriptome, sex was by far the most impactful and consistent parameter, overtaking diet and even strain differences (Figure 2A; Figure S2C). This was surprising because sex had comparatively the lowest effects on the phenotypes (Figure 1E), showing that strong sex-dimorphism at the RNA expression level does not necessarily translate to metabolic phenotypes. The second most impactful parameter was the mouse sub-species (Figure 2A), with the *Mus musculus musculus* (PWK/PhJ) and *Mus musculus castaneous* (CAST/EiJ) strains diverging from the other *Mus musculus domesticus* strains. Compared to the effects of sex and strain, the effects of diet were detectable only in strain- and sex-matched animals (Figure S2B), and only a handful of diet-driven genes were conserved across a limited number of strains. However, when we performed GSEA on the effects of diet or sex across strains, the situation was reversed, with many pathways showing a conserved response to diet across strains (Figure 3B; Figure 3C), while very few pathways were sex-dependent. In fact, the sex effects seem to be limited to a small number of sexually-dimorphic genes showing huge contrasts between sexes, that do not belong to a shared annotated pathway. The strongest sex-dimorphic genes were even conserved in humans, but their sex-specificity was sometimes inverted (Figure S3). Conversely, while the diet affects many different genes in different strains, those genes converge towards increased immunity-related genesets and a decrease in translation (Figure 3C). There are nevertheless exceptions, most notably the A/J strain females increasing translation-related genesets on HFD, again highlighting the importance of the genetic background.

The activity of the mitochondrial complexes in the liver was highly dependent on genetic effects rather than environmental factors (Figure 4C). In addition, the activity of mitochondrial complexes, particularly I and V, was positively correlated with indicators of physical fitness, such as maximal aerobic capacity (VO_2_ Max) and maximum run distance, indicating that optimal liver metabolic activity may be an important determinant of fitness (Figure 4E). These results suggest that specific ETC complexes measured in vivo or mitochondrial complex assembly may play a role in the liver response to dietary conditions, adding to the growing body of evidence for the central role of mitochondria in obesity and its complications^32^.

Our collection of phenotypic and molecular traits can be explored online. This represents a unique resource allowing researchers to link *in vivo* mitochondrial activity to phenotypic traits and transcriptomic data. To summarize the different parameters measured in this study, we show key phenotypic and molecular parameters associated with the response to HFD in each mouse strain (Figure 6). Each parameter was assessed relative to the other strains. Much like humans, each mouse strain responds differently to an obesogenic diet. The NZO/HILtJ strain had the more severe obesity and insulin resistance phenotype, yet transcript results of GSEA related to liver fibrosis and inflammation markers were comparatively lower. Conversely, the CAST/EiJ strain gained almost no weight and displayed no signs of glucose intolerance and insulin resistance, and yet it had some of the more pronounced transcript expressions of fibrotic and inflammation markers in the liver. While males in general displayed most severe symptoms, this was not the case in every strain, as female NOD/ShiLtJ mice were generally less affected by HFD than males. Thus, our overview of these strain-specific profiles (based on their phenotypic, transcriptomic and mitochondrial signatures), will guide researchers in choosing an appropriate model for correctly representing the different measurable outcomes in metabolic syndromes.

**Figure 6:**
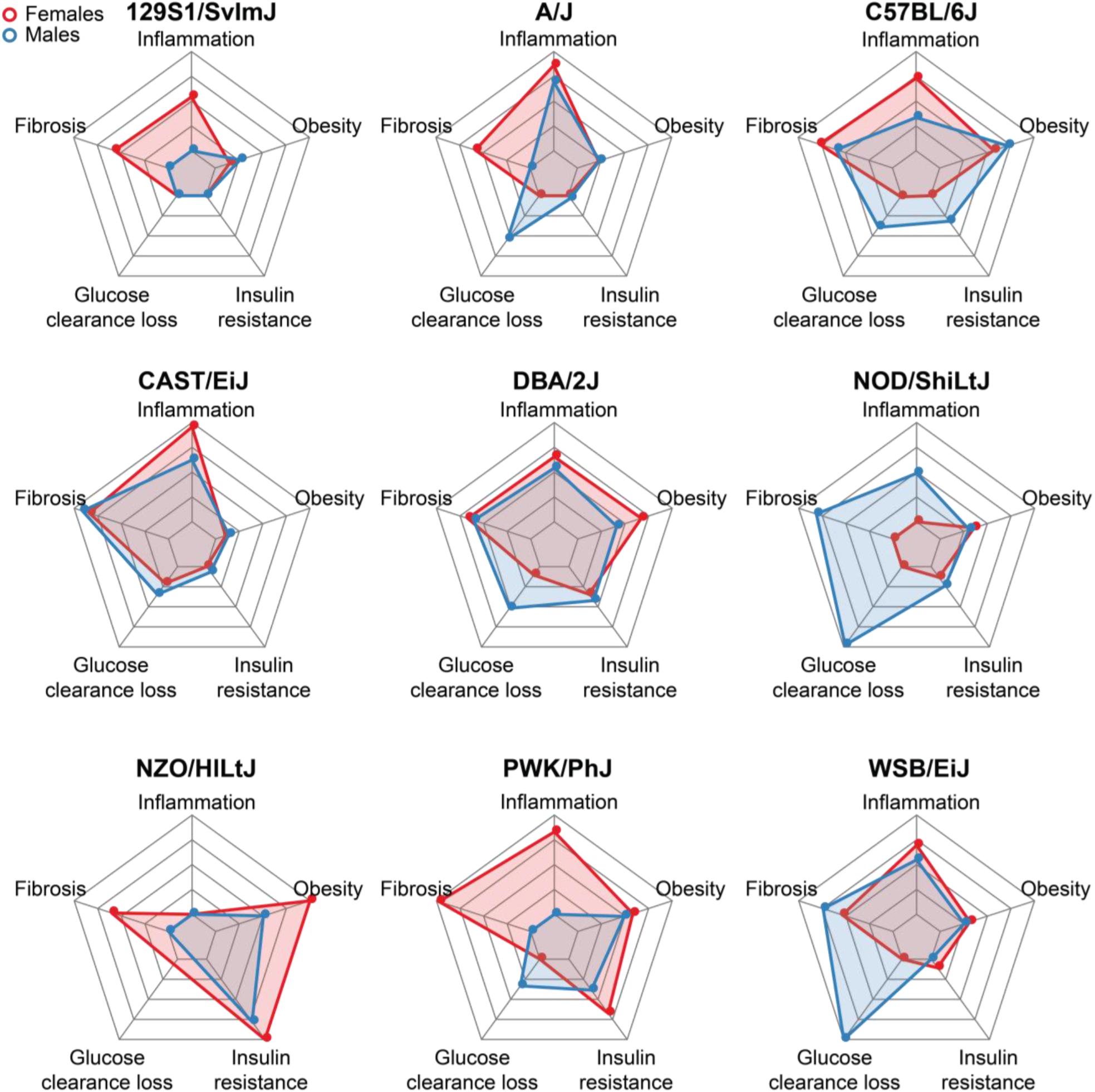
Strain-specific response signatures to obesogenic diet. Radar plots of the some hallmarks reflecting predisposition to metabolic disease in each strain. Each parameter is expressed as a percentage of the highest strain. Fibrosis and inflammation scores are based on the normalized enrichment score of the GSEA for “collagen-containing extracellular matrix” and “humoral immune response” respectively. Insulin resistance is based on homaIR value under HFD. Obesity is based on body weight gain relative to the starting weight before HFD. Finally, glucose clearance loss is the change in the slope of glucose clearance in the OGTT test upon HFD (missing data in the NZO strain, due to a too severe phenotype).

This study highlights the importance of considering biological factors like strains and genetic variations before drawing conclusions. By looking at varied strains of mice, some of which come from distinct subspecies or from the wild, we can approach the phenotypic variety present in human populations, and find conserved features, strengthening our conclusions. This is particularly important considering the number of discoveries that are affected by hidden biological and experimental parameters. Overall, our work presents the variation of *in vivo* mouse metabolic, transcriptomic and mitochondria-related traits and how they are affected by genetic background, sex and diet. We expect these data will be an important resource for the community that builds on previous studies of the collaborative cross founders where researchers measured other known or novel phenotypes and assessed biological covariates^15^ to enable the generation of new hypotheses in a data-driven manner.

## Materials and Methods

### Population handling

Mouse strains were imported from Charles River and bred at the École Polytechnique Fédérale de Lausanne (EPFL) animal facility for more than two generations before incorporation into the study. We examined 15 cohorts of 9 mouse inbred strains fed a CD or HFD, separated randomly into two groups of five mice of each sex for each diet. Strains were entered into the phenotyping program randomly and had staggered entry, typically by 1 or 2 weeks. HFD feeding started at 8 weeks of age. CD is Harlan 2018 (6% kCal of fat, 20% kCal of protein, and 74% kCal of carbohydrates), and HFD is Harlan 06414 (60% kCal of fat, 20% kCal of protein, and 20% kCal of carbohydrates). All mice were housed under 12 hours of light alternated with 12 hours of dark, with *ad libitum* access to food and water at all times. Body weight was measured weekly from 8 weeks of age until killing. All research was approved by the Swiss cantonal veterinary authorities of Vaud under license 2257.2. In accordance with the ethical license’s protocol, some animals were sacrificed early due to displaying signs of severe illness. Most notably, all but one of the NZO/HILtJ mice under HFD were sacrificed due to severe diabetes. In addition, the oral glucose tolerance test (OGTT) was omitted in the NZO/HILtJ mice, due to concerns about the stress of repeated blood collection.

### Phenotyping experiments

A visual summary of the phenotyping program is also included in (Figure 1A). At 14 weeks of age, after 8 weeks of dietary treatment, the cohorts underwent their first phenotyping test: 48 hours of respiration measurements in individual metabolic cages (Oxymax/CLAMS, Columbus Instruments). The first 24 hours were considered adaptation, and the second 24 hours were used for data analysis, including analysis of movement, the volume of oxygen inhaled, the volume of carbon dioxide exhaled, and derived parameters of these two, such as the respiratory exchange ratio (RER). Two weeks later, all cohorts underwent an oral glucose tolerance test. Mice were fasted overnight before the test, and fasted glucose was tested with a glucometer at the tail vein. All individuals were then weighed and given an oral gavage of 20% glucose solution at 10 mL per kg of weight. Glucometer strips were used at 15, 30, 45, 60, 90, 120, 150, and 180 min after the gavage to examine glucose response over time. Blood was also collected at 0 (pregavage), 15, and 30 min to examine insulin levels. Two weeks later, at 18 weeks of age, we performed a cold response test. The basal body temperatures of mice were examined rectally, after which mice were placed individually in prechilled cages in a room at 4°C. The cages were the standard housing cages but with only simple wood chip bedding. Body temperature was checked every hour for 6 hours, after which the mice were returned to their normal housing cage. Two weeks later, at 19 weeks of age, the mice were placed individually in regular housing cages for basal activity recording. The housing cages were then placed in laser detection grids developed by TSE Systems (Bad Homburg, Germany). Within the cages, woodchip bedding was retained. Food and water were as normal throughout the standard housing, both of which require rearing to reach. The detection grid has two layers: one for detecting horizontal (X-Y) movement (“ambulation”) the other for vertical (Z) movement (“rearing”). Mice were housed individually for the 48-hour experiment starting at about 10 a.m., with the night cycles (7 p.m. to 7 a.m., with 30 min of both dawn and dusk) used for movement calculations. After this 2-day experiment, all mice performed a VO2max treadmill experiment using the Metabolic Modular Treadmill (Columbus Instruments). During the first 15 min in the machine for each individual, the treadmill was off while basal respiratory parameters were calculated. The last 2 minutes of data before the treadmill turned on are considered basal levels (most mice spend the first few minutes exploring the device). The treadmill then started at a pace of 4.8 m per minute (m/min), followed by a gradual increase over 60 s to 9 m/min, then 4 min at that pace before increasing to 12 m/min over 60 s, then four min at that pace before increasing to 15 m/min over 60 s, then 4 min at that pace, then the speed increased continuously by 0.015 m per second^2^ (or +0.9 m/min^2^) thereafter until the end of the experiment at 63.5 min, 1354.5 m, or when the mouse is exhausted. CD cohorts ran against a 10° incline, whereas HFD cohorts were set at 0°. For this test, all mice were taken out when exhausted or when physical constraints were too high. The distance, maximum VO2, and maximum RER were recorded. Maximums must be consistent across multiple measurements, and not single-measurement spikes, which were removed. At 21 weeks of age, mice were fasted overnight before they were killed. In addition to the body weight measurements taken each week and before each phenotyping experiment, body composition was recorded at week 14. To do so, each mouse was placed briefly in an EchoMRI (magnetic resonance imaging) machine (the 3-in-1, Echo Medical Systems), where lean and fat mass are recorded, along with total body weight, taking ∼1 min per individual. Lean mass is used as a corrective factor for respiratory calculations from the Comprehensive Lab Animal Monitoring System (CLAMS). All other tests are normalized to total body weight in our analyses.

### Sample collection

The sacrifices took place from 8:30 a.m. until 10:30 a.m., with isoflurane anesthesia followed by a complete blood draw (∼1 mL) from the vena cava, followed by perfusion with phosphate-buffered saline. Half of the blood was placed into lithium-heparin (LiHep)–coated tubes and the other half in EDTA-coated tubes; then both were shaken and stored on ice, followed immediately by collection of the liver. The LiHep blood taken for plasma analysis was also centrifuged at 4500 revolutions per minute (rpm) for 10 min at 4°C before being flash-frozen in liquid nitrogen. Whole blood taken for cellular analysis was processed immediately after the killing (i.e., after ∼1 to 2 hours on ice). Gallbladders were removed, and the livers were cut into small pieces before freezing in liquid nitrogen until preparation into mRNA, protein, or metabolite samples. Liver and blood serum were then stored at –80°C until analysis.

### Sample preparation and analysis

For mRNA, liver tissue were crushed in liquid nitrogen and then 10 mg of tissues were suspended in TRIzol (Invitrogen) and homogenized with stainless steel beads using a TissueLyser II (Qiagen) at 30 Hz for 2 min. RNA was extracted and purified using Direct-zol-96 RNA kits (Zymo Research). mRNA concentration was measured for all samples. All samples passed a quality check of purity (NanoDrop) and fragmentation (FragmentAnalyzer).

### RNA-sequencing and mapping

The STAR aligner^33^ was used for mapping the RNA-Seq data to the C57BL/6J reference genome and determining gene counts. We did not use distinct genomes for each strain due to various genome quality differences between mouse strains that could create bigger artefacts than mapping all strains on the same reference genome in terms of mapping efficiency and gene count estimation. Differential gene expression was done using the limma R package and the voom method^34^. Gene set enrichment analysis (GSEA) was done using the GSEA method of the clusterprofiler R package^35^. Genes were ranked according the signed –log10(BH adjusted p-value) obtained by looking at the diet effect in female or male in each strain separately. We selected the genesets from the biological process of the gene ontology which had the highest significance levels in the overall response to the diet in each sex.

### Mitochondrial activity

Measurement of mitochondrial activity were performed by Metabiolab France. The method is based on spectrophotometry and mitochondrial activity of the different complexes of the oxidative phosphorylation chain was normalized by citrate synthase activity (when not stated otherwise).

### Human data analysis

We have access to the Genotype-Tissue Expression (GTEx) project^36^version 8 through accession number dbGaP phs000424.v8.p2. When assessing the effect of sex on gene expression in GTEx livers, we aimed to correct for unmeasured covariates and hidden factors trough the addition of Probabilistic Estimation of Expression Residuals (PEER) factors. To analyze sex and age, we re-calculated the PEER factors following the pipeline from the GTEx consortium^37^, with two modifications. First, removed mitochondrial-genes, as well as X-and Y-chromosome-encoded genes. Second, we included the first five genotyping principal components, the sequencing protocol (PCR-based or PCR-free), the sequencing platform (Illumina HiSeq 2000 or HiSeq X), sex, age, quadratic age, the interaction between sex and age, and the interaction between sex and quadratic age in the new PEER factors. We corrected for sequencing depth, gene length and GC percentage with through conditional quantile normalization with the cqn R package^38^ and estimated dispersion parameters with DESeq2’s estimateDispersions function^39^. In addition to the variables mentioned in the PEER factor calculations, we also included the 20 first re-calculated PEER factors as predictors in DESeq2’s negative binomial generalized linear models. While the ages of the GTEx subjects range from 21 to 70 years old, all CC founder mice in this study were 21 weeks old at the time of sacrifice. To maximize the similarity between the mouse and human datasets, we used Wald tests to assess the average effect of sex on gene expression for a 30 years old subject. P-values were adjusted for multiple testing through DESeq2’s independent filtering procedure coupled to the Benjamini-Hochberg false discovery rate (FDR) correction with a significance cut-off at 5% FDR.

### Bioinformatic and statistical analysis

All the bioinformatics and statistical analysis were performed in R 3.5.2 and Rstudio Pro. Effect size was computed as follows: difference of the means divided by the global standard deviation. Correlation are based on Spearman’s correlation (except when stated otherwise). When needed, p-values were corrected for multiple testing with the Benjamin-Hochberg false discovery rate. We used the type II sum of squares, which shows the variance explained by a parameter conditional on the presence of all other main effects in the model. The type II sum of squares preserve the principle of marginality and are independent of the listing order of the model parameters. variantPartition package was used to compute the variance explained by defined parameters for transcriptomic data.

### Interactive data visualization and availability

All metabolic traits, mitochondrial activity and transcriptome data collected in this study can be explored with an online, interactive interface at https://lms.shinyapps.io/Project_CC_Founder_App/. The website is separated into 5 parts (description of the experimental design, the metadata, the phenotypic traits, the transcriptomic data and the mitochondrial activity). In addition to the exploration of the raw data it is possible to see the trait-trait correlations as well as the effect of diet and sex on each strain separately. At the transcript level, individual genes and genesets can be visualized as well as the effects of the diet and sex in any subset of strains. The raw transcriptomic data has been submitted to GEO expression database under the accession <u>GSE182668</u>. The individual phenotype data and mitochondrial data is provided on table S1.

## Acknowledgments

We thank the members of J. Auwerx’s laboratory for their help with sacrifices and sample processing and the staff of EPFL’s Center of Phenogenomics (CPG) for help with phenotyping. We thank all members of J. Auwerx and K. Schoonjans laboratories for helpful discussions. We thank Metabiolab for their help in measuring the activity of mitochondrial complexes.

## Author contributions

A.M.B., M.B.S and J.A. conceived the project. A.M.B., S.R., G.B. and T.I.L. performed the experiments. A.M.B., J-D.M., L.J.E.G. performed data analysis. G.E.A. worked on the online website. J.D.M., M.B.S., and J.A. supervised the study. A.M.B., J-D.M. and J.A. wrote the manuscript with comments from all authors.

## Data and materials availability

The authors declare that all the data supporting the findings of this study are <u>available within the paper and its supplementary materials</u>.

## Supplementary figures

**Figure S1:**
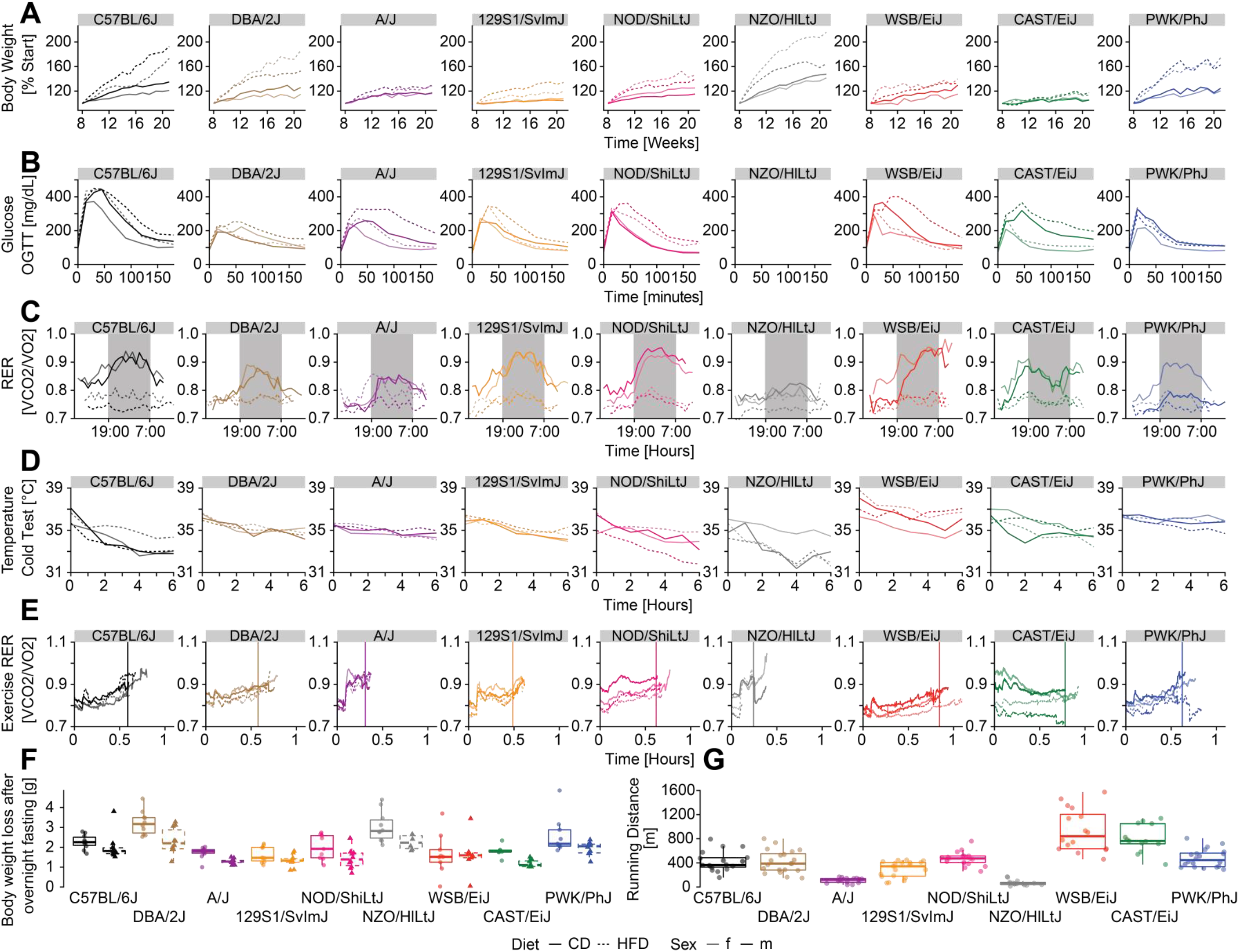
Variation of metabolic traits collected in this study. A) Body Weight (expressed as percentage from the individual starting bodyweight before HFD). B) Glucose levels during the course of an oral glucose gavage test (OGTT). C) Circadian respiratory exchange ratio (RER) over a day using a metabolic chamber. D) Rectal temperature taken during a cold test. E) Uphill exercise respiratory exchange ratio (RER) during a VO_2_max experiment. Vertical line represents the mean time run per strain. F) Body weight loss after an overnight fasting. G) Distance run by the mice during the VO_2_max experiment.

**Figure S2:**
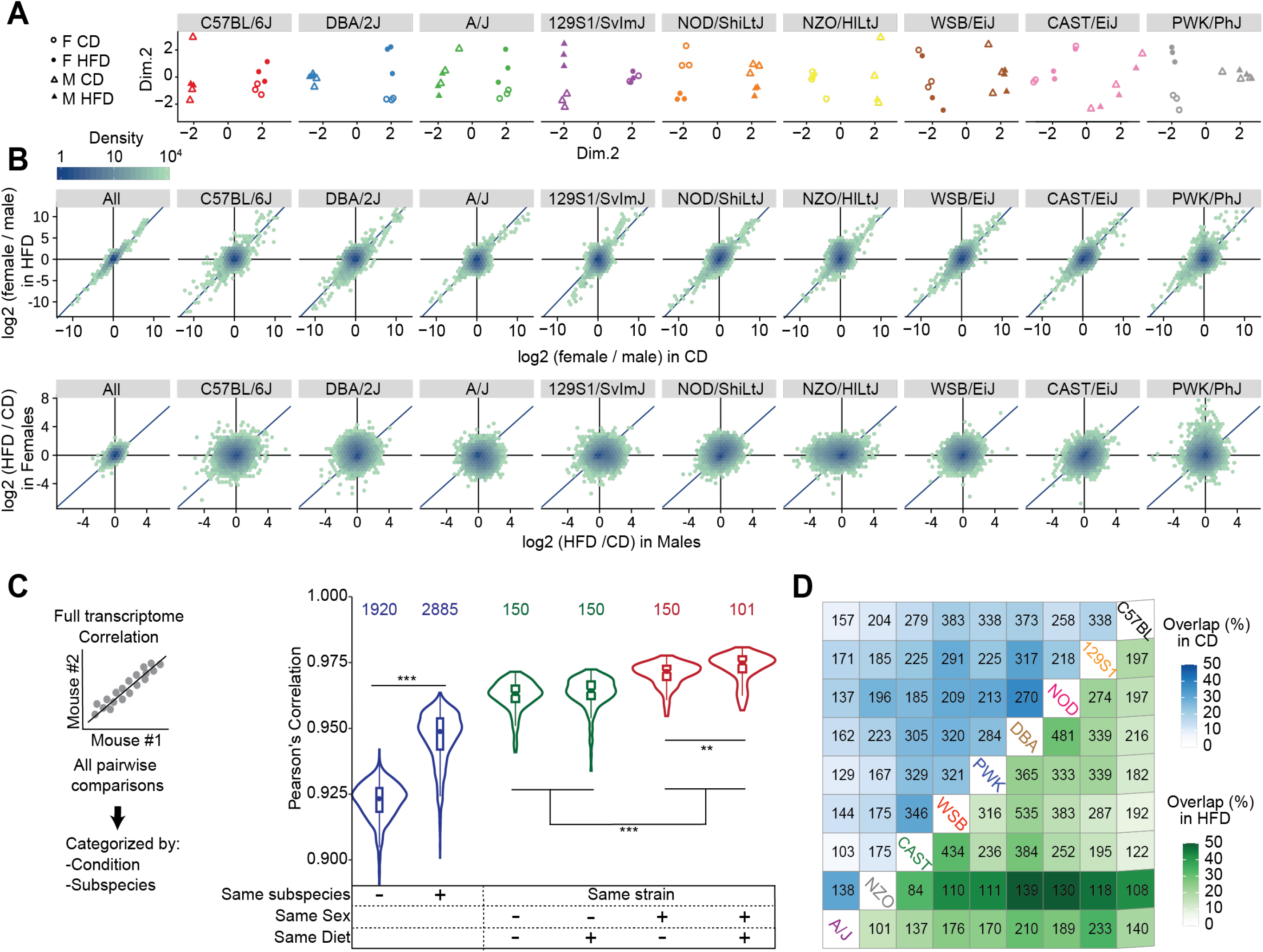
Strain-specific liver gene expression response to diet of is not well conserved across sexes. A) Multidimensional scaling (MDS) for transcriptomic data. Euclidean distances between each strain were calculated pairwise based on the leading log_2_-fold changes for the 500 most differentially expressed genes. Each dot represents a mouse in a given condition, which is in turn associated with a specific shape. The effect of sex is predominant compared to the effect of the diet, which can be seen in some strains for one or both sexes. B Top: overall and per-strain correlation plots of sex effects (log_2_ fold changes of female vs male) in CD vs HFD. Bottom: overall and per-strain correlation plots of diet effects (log_2_ fold changes of HFD vs CD) in females vs. males. Sex differences are well conserved between CD and HFD as well as between strains. Diet differences are qualitatively and quantitatively poorly conserved across sexes and strains. Sex differences are well conserved between CD and HFD as well as between strains. Diet differences are qualitatively and quantitatively poorly conserved across sexes and strains. C) Pearson’s correlation of pairwise mice comparisons of global gene expression. Note a strong effect of subspecies differences on transcriptomic profiles. The effects of sex and diet are subordinate to the subspecies effect. D) Number (written) and percentage (color legend) of overlapping genes that are differentially expressed between females and males, in CD (blue) or HFD (green).

**Figure S3:**
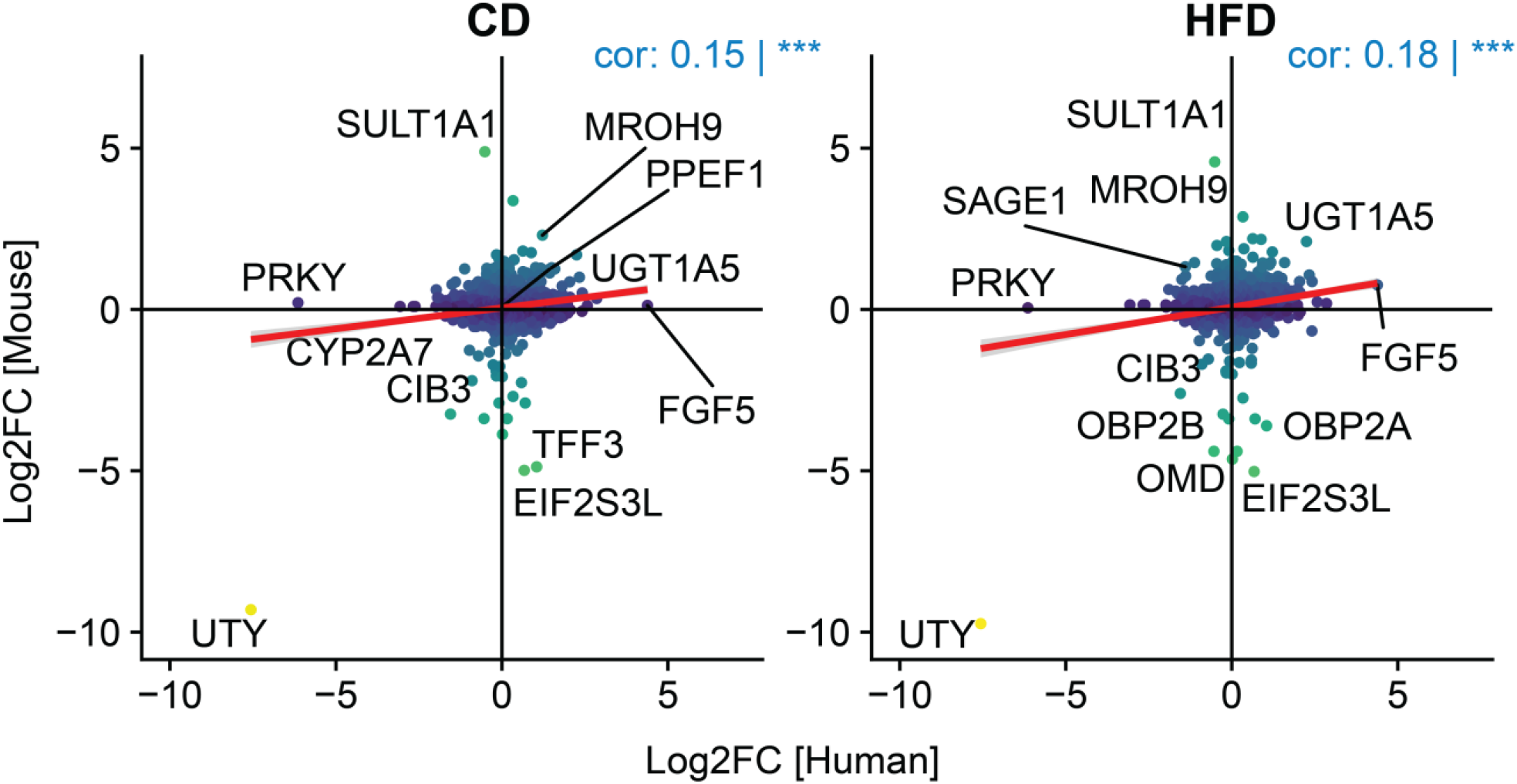
Correlations of the fold change between sexes in mouse and human liver transcriptomes. The human liver transcriptome was extracted from the GTEx v8 database. Colour scale is based on the fold change in mice.

